# NECo: A node embedding algorithm for multiplex heterogeneous networks

**DOI:** 10.1101/2020.06.15.149559

**Authors:** Cagatay Dursun, Jennifer R. Smith, G. Thomas Hayman, Anne E. Kwitek, Serdar Bozdag

## Abstract

Complex diseases such as hypertension, cancer, and diabetes cause nearly 70% of the deaths in the U.S. and involve multiple genes and their interactions with environmental factors. Therefore, identification of genetic factors to understand and decrease the morbidity and mortality from complex diseases is an important and challenging task. With the generation of an unprecedented amount of multi-omics datasets, network-based methods have become popular to represent the multilayered complex molecular interactions. Particularly node embeddings, the low-dimensional representations of nodes in a network are utilized for gene function prediction. Integrated network analysis of multi-omics data alleviates the issues related to missing data and lack of context-specific datasets. Most of the node embedding methods, however, are unable to integrate multiple types of datasets from genes and phenotypes. To address this limitation, we developed a node embedding algorithm called Node Embeddings of Complex networks (NECo) that can utilize multilayered heterogeneous networks of genes and phenotypes. We evaluated the performance of NECo using genotypic and phenotypic datasets from rat (*Rattus norvegicus*) disease models to classify hypertension disease-related genes. Our method significantly outperformed the state-of-the-art node embedding methods, with AUC of 94.97% compared 85.98% in the second-best performer, and predicted genes not previously implicated in hypertension.

**Availability and implementation:** The source code is available on GitHub at https://github.com/bozdaglab/NECo.

## I. Introduction

Almost two-thirds of the deaths in the U.S. are caused by complex diseases such as cardiovascular disease, hypertension, cancer and diabetes [9]. Complex diseases involve interactions of multiple genes with each other and with environmental factors [23]. Identification of complex disease genes to understand disease pathways and mechanisms, and to reduce death rates, is therefore a vital yet difficult endeavor. Comprehensive understanding of the mechanisms of complex diseases and traits often involve generating large datasets that characterize the effects on phenotypes by, for instance, the genome, transcriptome, epigenome or proteome. These multiomics datasets, however, might have patterns of missing data across the different datasets and often the available datasets are not specific for the particular complex trait in question, such as protein-protein interactions (PPI) [36]. To facilitate better understanding of the complexity of multilayered molecular interactions and elucidate the genotype-phenotype relations, integrative network analysis methods have been employed [5, 6, 15, 37].

To utilize such networks for downstream supervised and unsupervised analyses such as link prediction, community detection and node classification, latent representation of networks (i.e., node embeddings) are computed [12, 25]. Several approaches to learn latent representation of networks have been implemented and applied in different domains [12, 24, 25, 34, 39, 40]. Many of the recent studies utilize simple networks for latent representation, and recently several tools have emerged to learn the latent representation of more complex networks [1, 7, 38, 40].

Some of the recent node embedding methods rely on random walks-based approaches to learn the neighborhood of the nodes in the network. In a random walk, an imaginary walker starts at an initial node and iteratively visits one of its neighbors in the network. Random walk with restart (RWR) is a variation of random walk where in the first step, an imaginary walker starting from an initial node moves to one of its immediate neighbors in the network and then either walks to another neighbor or jumps back to the initial node iteratively. As random walks use the topological structure of networks, they are effective to capture the proximity of nodes to each other, and they are scalable for large networks [10].

After learning the node neighborhoods, node embeddings are computed by using approaches inspired from the word2vec algorithm, which is a word embedding algorithm [22]. Given a text, word2vec uses the Skip-gram algorithm to learn the word embeddings. The Skip-gram algorithm predicts surrounding words of a given word based on the assumption that the words appearing more frequently in the same context are similar to each other [25].

DeepWalk is a random walk-based node embedding algorithm which utilizes homogeneous networks [25]. It generates node sequences using truncated random walks then utilizes the Skip-gram algorithm to learn the node embeddings. DeepWalk employs a hierarchical softmax as a normalization factor to speed up the process of the Skip-gram. Node2vec is another random walk-based node embedding algorithm for homogeneous networks [12]. There are two key differences between Node2vec and DeepWalk; first, node2vec uses negative sampling as a normalization factor; second, it uses a biased random walk to capture both structural similarities of nodes as well as homophily [13]. Graph Diffusion Convolution (GDC), also for homogenous networks, differs from previous random walk-based node embeddings by transforming the underlying network structure using heat kernel or RWR [16]. Metapath2vec, another random walk-based node embedding algorithm, was developed to address the need for heterogeneous (having multiple node types) networks [7]. It has two key features; it utilizes meta-paths to guide the random walks and it can apply heterogeneous negative sampling based on the number of node types in the heterogeneous network. OhmNet utilizes hierarchical multilayer network structure of proteins from different tissues to learn the latent representation of proteins [40]. Multiplex network embedding (MNE) was developed to utilize multiplex (multiple layered) network structure and to generate a common embedding vector across all layers of the network and specific embedding vectors for each layer [38]. Multi-net utilizes the multiplex network structure by taking into a uniform transition probability for jumping to other layers [1]. GATNE was developed to utilize multiplex heterogeneous networks while integrating node attributes. It has two versions: GATNE-T is using network topology and GATNE-I is using attributes of nodes in addition to the network topology based on truncated random walk strategy [4]. It uses meta-paths to handle heterogeneous networks. However, these methods lack the ability to utilize multiplex heterogeneous networks using steady state ranking of RWR, and they cannot efficiently utilize the proximity of heterogeneous nodes.

In this study, we present a new node embedding algorithm, called Node Embeddings of Complex networks (NECo), that can utilize multiplex heterogeneous networks. NECo utilizes multiplex gene and phenotype networks to learn the latent features of genes and phenotypes for downstream analysis such as gene function classification and drug-gene interactions (Fig. 1). First, NECo creates a complex network structure using multiple gene/phenotype layers and a bipartite network of genes and phenotypes. NECo uses a RWR strategy to generate the top *N* node neighborhoods for each node in the network. Unlike Metapath2vec NECo handles heterogeneous nodes by generating four types of neighborhoods: i) gene-gene: gene rankings starting from each gene node, ii) gene-phenotype: phenotype rankings starting from each gene node, iii) phenotype-gene: gene rankings starting from each phenotype node, iv) phenotype-phenotype: phenotype rankings starting from each phenotype node. Then, it learns the embeddings of genes and phenotypes based on *top N* nodes of each of these neighborhoods utilizing the Skip-gram algorithm. For gene embeddings gene-gene, gene-phenotype and phenotype-gene neighborhoods are used. For phenotype embeddings phenotype-phenotype, phenotype-gene and gene-phenotype neighborhoods are used. Finally, those different node embeddings are concatenated and utilized by supervised/unsupervised learning algorithms for various downstream analyses.

**Fig. 1.**
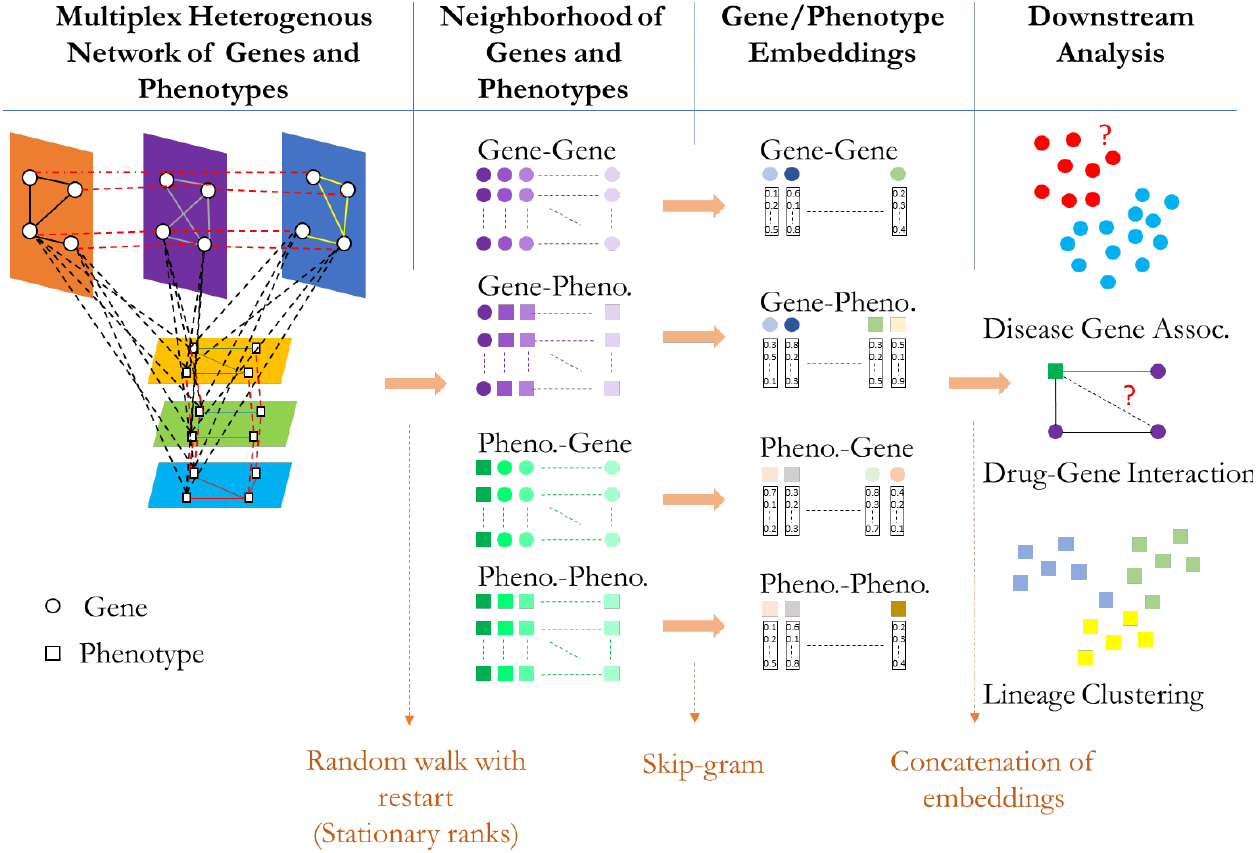
Overview of the NECo framework. NECo first generates the complex multiplex heterogeneous undirected weighted network. Then, it obtains the neighborhood of Gene-Gene, Gene-Phenotype, Phenotype-Gene and Phenotype-Phenotype by random walk with restart. Thirdly, NECo takes the *top N* of those neighborhoods and learns the latent representation of nodes using the Skip-gram algorithm. The learned node embeddings of different spaces then are concatenated and used for classification using a statistical learning algorithm.

We compared NECo’s performance with other approaches on predicting known hypertension disease-related genes using multidimensional rat datasets. NECo outperformed the other approaches by about a 9% margin. Furthermore, the top 20 novel hypertension-related gene predictions by NECo had supporting evidence in the literature of their role in hypertension.

## II. MATERIALS AND METHODS

### A. Random Walk on Multiplex Heterogeneous Network

NECo utilizes RWR algorithm on undirected multiplex heterogeneous networks to compute a node neighborhood starting from each node in the network and applies the Skipgram algorithm to learn the latent features of nodes in the network based on their RWR rankings (Fig. 1). NECo creates a multiplex heterogeneous network of genes and phenotypes using PhenoGeneRanker with default parameters [8]. Specifically, NECo can utilize multiple layered undirected networks which have two different type of nodes. Unlike several node embedding algorithms [7, 12, 25] that rely on simulated random walks, NECo utilizes the steady state distribution of RWR to generate the neighborhood of the nodes in the network (Eq. (1)),

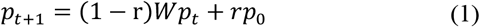

where *r* is the restart probability (i.e., RWR moves backs to the starting nodes), *p*_*t*_ represents the probability distribution vector of nodes at time *t* and *W* is the transition matrix of the network (Eq. (2)), which is calculated by the multiplication of degree diagonal matrix (*D*) of the network and the adjacency matrix of the network (*A*).

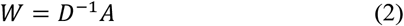

After a number of steps, Eq. (1) reaches a steady state for undirected networks [6]. The magnitude of *r* affects the convergence rate of the RWR algorithm, where a large *r* leads to fast convergence to steady state [18] and limits the diffusion of the random walk. The steady state distribution (*p*_*s*_) can be used as a proximity vector for the nodes in the network starting from an initial node. NECo sets *r* = 0.7 by default as in other RWR algorithms [17, 20, 29, 33].

### B. Node Embedding

NECo utilizes *top N* nodes of different RWR neighborhood spaces as a proximity measure of the node sequences (Fig. 1). NECo generates mixed node rankings that include genes and phenotypes starting from either a gene or phenotype node via RWR. To generate Gene-Gene and Gene-Phenotype neighborhoods, NECo separates the node rankings by node type where the initial node for RWR is a gene node, and it generates Phenotype-Gene and Phenotype-Phenotype neighborhoods similarly where the initial node is a phenotype node. Then, NECo utilizes the Skip-gram algorithm to learn the node embeddings based on the neighborhoods of each gene or phenotype. The goal of the Skip-gram algorithm is to learn the node features of a given node which are predictive of nodes in its neighborhood. The objective function of the Skip-gram algorithm becomes the maximization of the log likelihood in Eq. (3);

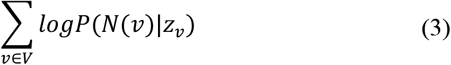

where *V* is the set of nodes in the network, *ν* is a node in the network, and *N*(*ν*) is the set of neighbor nodes of node *ν*, and *Z*_*v*_ is the embedding of node *ν*. Symmetricity assumption of the learned features leads to the softmax function in Eq. (4) that gives the probability of obtaining node *u* given the node *ν* in a given graph.

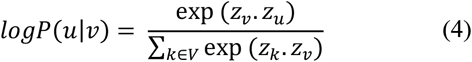

We use negative sampling for efficient calculation of the softmax in Eq. (4) [21].

For this study, the aim of the generated node embeddings was to predict the hypertension disease related genes; therefore, NECo was not used to generate phenotype embeddings. However, it is straightforward to generate phenotype embeddings for a different problem setting such as drug-gene interaction studies.

### C. Complex Network of Rat Datasets

To evaluate the performance of NECo, we applied it to a multidimensional rat dataset to generate a multiplex heterogeneous network, with the goal of predicting hypertension disease-related rat genes. We created a three-layer gene interaction network, namely gene transcript co-expression, protein-protein interaction and pathway layers. We then created a three-layer phenotype network using rat strain annotations; mammalian phenotype ontology (MPO) term-based similarity; disease ontology (DO) term-based similarity; quantitative phenotype (QP) measurements-based similarity of rat strains. We connected the gene multiplex network to the phenotype network based on the MPO annotation-based similarity of genes and strains. All the layers were composed of undirected and weighted edges. NECo learns the node embeddings utilizing this multiplex heterogeneous network.

1. *Gene Network*: We created a multiplex gene network using PPI, pathway and co-expression layers.

a. *PPI Layer*: To create the PPI layer, all physical interactome data for *Rattus norvegicus* were downloaded from the STRING V11 database [32]. For protein pairs having multiple interactions, we aggregated their interactions by taking the arithmetic mean of the interaction weights. Protein IDs were mapped to gene IDs by using an alias file in the STRING database. Proteins that were mapped to the same genes were merged into a single node, and the average of their interaction weights were used as the merged interaction weight.
b. *Pathway Layer:* To create the pathway layer, *Rattus norvegicus* pathway annotation of genes and the pathway ontology tree were downloaded from the Rat Genome Database (RGD) [19]. Semantic similarity of genes was calculated using the ontologyX R package [11]. Ontology-based semantic similarity measures the degree of relatedness between two entities according to the similarity in meaning of their annotations over a set of ontology terms [27]. We applied a best-match-average approach for two sets of terms which is given by the average similarity between each term in the first set and its most similar term in the second set averaged with its reciprocal [26]. To increase the computational efficiency and decrease the number of uninformative correlations, a maximum possible hard threshold value of 0.5668 was applied to shrink the number of edges while keeping the network connected.
c. *Co-expression Layer*: The co-expression layer was based on our RNA-seq expression dataset (GSE50027) [35] of liver tissue from six Lyon Hypertensive (LH) and six Lyon Normotensive (LN) rats obtained from the Gene Expression Omnibus (GEO) database [2]. We applied a filtering metric of zFPKM [14] to filter the active genes from the background genes in the dataset. We used zFPKM > −3 to select expressed genes as recommended in [14]. We removed the genes having undefined values for four or more samples after zFPKM calculation. We calculated the co-expression weights of remaining genes for normotensive and hypertensive samples using the Pearson correlation. We then calculated the differential co-expression weight of genes by calculating the natural log ratio of co-expression weights between hypertensive and normotensive samples. Finally, we applied a maximum possible hard threshold value of 7.2393 to shrink the number of edges while keeping the network connected.
2. *Strain Network*: We created a multiplex phenotype network using MPO, DO and QP measurements-based strain similarity layers.

a. *MPO- and DO-Based Strain Similarity Layer*s: The MPO- and DO-based strain similarity layers represent the strain similarity based on MPO and DO annotations of strains obtained from RGD [31], respectively. To make these layers more context-specific, we computed similarity scores based on hypertension-related MPO/DO terms. First, we determined a list of hypertension related MPO/DO terms (Supplementary Table 1 and 2) and calculated vectors that represent each strain based on their MPO/DO annotation semantic similarity to the hypertension related MPO/DO term vector. Second, we calculated the similarity of strains as the dot-product of those semantic similarity vectors. Finally, we applied a maximum possible hard threshold value of 0.1032 for MPO-based and 0.0740 for DO-based strain similarity networks to minimize the number of edges while keeping the networks connected.
b. *QP Measurements-Based Strain Similarity Layer*: To create the QP layer, we downloaded the quantitative phenotype measurements namely systolic blood pressure, heart rate and heart weight annotated to strains in RGD. To prevent measurement bias between different sexes, we used the studies having male samples only as the number of female samples were very low. To consider measurements from adult rats only, we set the range of the age of samples to be 28 to 410 days. We utilized only *in viv*o measurements for systolic blood pressure to prevent measurement bias. We calculated Euclidean distance of the strains using these three measurements. We took the multiplicative inverse of the distance and applied a maximum possible hard threshold value of 6.1 to minimize the number of edges while keeping the network connected.
3. *Gene-Strain Bipartite Layer*: The gene-strain bipartite layer represents the relatedness of genes to strains and connects the gene nodes to the strain nodes based on their semantic similarity of MPO annotations. We downloaded the gene MPO annotations from RGD. To increase the number of covered genes in the bipartite layer, we utilized MPO annotations of mouse orthologs of rat genes by downloading mouse MPO annotations from Mouse Genome Informatics [30]. We utilized RGD MPO annotation for strains. Then, we calculated the semantic similarity following a strategy similar to the gene pathway layer.

### D. Disease Gene Classification Using Gene Embeddings

To evaluate the performance of NECo embeddings we used multiple gene and phenotype datasets of rat to predict hypertension disease related rat genes. We ran NECo on different network configurations. We omitted single gene/phenotype layers to compare the contribution of different layers. We also created multiplex and aggregated gene/phenotype networks to compare the contribution of multiplex gene/phenotype networks. We created aggregated networks by taking the union of the nodes and calculated the geometric mean of edge weights if the edges were common in the aggregated layers. We employed the Generalized Linear Model (GLM) to classify hypertension-related genes using the gene embeddings computed by NECo. The feature set of each gene was composed of concatenation of gene embeddings based on Gene-Gene and Gene-Phenotype neighborhood spaces. We did not use gene embeddings based on Phenotype-Gene neighborhood space as they did not improve the prediction accuracy when concatenated with other gene embeddings. We used the rat gene disease annotations in RGD to determine the set of “ground truth” hypertension disease-related rat genes. We selected the experimental annotations but excluded the genes having only expression based experimental evidence codes (Supplementary Table 3). The number of unique genes in the whole network where all gene layers were used was 18,275 and the number of genes in the “ground truth” set was 167. We employed 10-fold 10-repeat stratified cross validation for performance measurement.

### E. Comparison with Existing Node Embedding Algorithms

We compared NECo with DeepWalk [25], Node2vec [12] and Metapath2vec [7] node embedding algorithms.

For DeepWalk, we used the node2vec implementation as it is stated that the performance difference between hierarchical softmax and negative sampling normalizations in the Skip-gram algorithm is negligible [7]. The hyperparameters *p* and *q* of Node2vec control the biased random walk; *p* controls the likelihood of returning to the previously visited nodes, while *q* controls the likelihood of visiting further nodes. We set *p* = *q* = 1 for an unbiased walk for DeepWalk. Since DeepWalk can only utilize a homogeneous network, we generated an aggregated gene network of three gene layers by taking the geometric mean of the edge weights and keeping all the genes in the network. DeepWalk has several hyperparameters; the *number of walks* controls the number of runs of random walk per node, *walk length* controls the number of time that random walker will hop per random walk, *context size* controls the size of *window*, which defines the group of co-occurring nodes in the Skip-gram algorithm, *embedding size* controls the latent feature vector of a node that will be learned by the Skip-gram algorithm. We ran DeepWalk for all combinations of key hyperparameter value choices (i.e., *number of walks per node*: {10, 20}, *walk length*: {80, 100}, *context size*: {5, 10} and *embedding size*: {128, 256, 512}).

We utilized the same grid search for the common hyperparameters of node2vec and DeepWalk. We also input the same aggregated gene network to Node2vec. For *p* and *q* values we used grid search for the interval of {0.25, 0.5, 1, 2}.

For evaluating the performance of Metapath2vec, we used bipartite relations between genes and strains, and guided the random walks of Metapath2vec using the gene-strain-gene metapath scheme. We input the unweighted gene-strain bipartite network to the Metapath2vec algorithm as it can only utilize unweighted bipartite networks. Number of walks was set to 1000, and walk length was set to 100 as they were best performing values in [7]. The same embedding size values were used with DeepWalk and Node2vec, *context size* had values of {5, 7, 10, 15} following [7]. A grid search was applied for these hyperparameters.

## III. Results

In this study, we developed a node embedding algorithm, NECo that uses multiplex gene and phenotype networks to learn node embeddings. Using a multidimensional rat dataset, a multiplex gene network was generated using differential transcript co-expression, PPI and pathway layers. A multiplex phenotype network was generated using MPO, DO and QP strain layers. We applied a grid search for different hyperparameters (e.g., *top N*, *embedding size*, *context size*) for each embedding space across different network combinations (See II.C).

We present our findings on comparing NECo to other state-of-the-art node embedding algorithms and show literature review results for the novel predictions by NECo. We also present results on hyperparameter analysis and the effects of different embedding spaces, multiplex networks compared to aggregated networks, and of individual network layers.

### A. NECo Outperforms Other Node Embedding Algorithms

We compared NECo with three state-of-the-art node embedding algorithms, namely Node2vec, DeepWalk and Metapath2vec. We built a complex network based on rat datasets and computed embeddings for each gene using the node embedding algorithms. Then we evaluated the performance of hypertension disease-related gene prediction (See Section II.C and II.D).

Table 1 shows mean area under the receiver operating characteristic curve (AUC), F1 micro and macro scores of classification using the gene embedding of each algorithm. GLM classification based on the embeddings computed by NECo achieved an AUC of 94.97% using a multiplex gene network of three layers and an aggregated phenotype network of MPO and QP layers, whereas the second-best performer classification, which was based on the embeddings computed by Node2vec was 85.98%. Classifications using NECo embeddings had higher F1 micro and macro scores than other methods, too.

**Table 1.** Mean area under receiver operating characteristic curve (AUC), F1 Micro and F1 Macro values with standard deviations are given for 10 runs for each embedding. For the GLM classification 10-fold cross-validation was repeated 10 times. Values are in percentage.

| Method | Mean AUC | Mean F1 Micro | Mean F1 Macro |
| --- | --- | --- | --- |
| NECo | 94.97 ± 0.24 | 90.98 ± 0.15 | 55.06 ± 0.13 |
| Node2vec | 85.98 ± 0.54 | 87.92 ± 0.28 | 51.48 ± 0.20 |
| DeepWalk | 85.21 ± 0.85 | 88.24 ± 0.27 | 51.88 ± 0.28 |
| Metapath2vec | 82.33 ± 0.05 | 51.12 ± 2.08 | 36.19 ± 1.01 |

### B. NECo Predicts Novel Hypertension-Related Genes

To further examine top false positive predictions by NECo, we performed a literature-based search of these predictions. To identify the top-ranked predicted genes, we picked the top-scored seven configurations based on AUC scores, as their AUC scores were nearly identical to each other (Supplementary Table 4). For each network configuration, we generated node embeddings 10 times and ranked the genes by their prediction probability based on the 10-fold 10-repeat stratified cross validation. We then calculated the final rank based on the median ranks of each gene across these 70 results. We chose median ranks instead of mean ranks to avoid any outlier ranks. We filtered the known hypertension genes based on our ground truth set of hypertension disease-related genes. We investigated the top 20 novel predictions for genes that at the time had not been annotated for hypertension-related disease at RGD (Table 2). Literature curation of these genes determined that 18 of NECo’s top predictions have published evidence, in most cases not merely expression-based, that the genes are involved in hypertension (Table 2). For the remaining two genes, we could not find specific hypertension-related annotations. However there is supporting literature that *Pla2g10* is involved in a signaling axis regulating blood pressure homeostasis [3] and *Ckmt2* is involved in cardiovascular disease [28].

**Table 2.** Novel neco gene predictions sorted by prediction probability with supporting publications. references for publications listed in supplementary Table 5.

| Gene Symbol | # of Supporting Publications |
| --- | --- |
| *Pdc* | 3 |
| *Rcn2* | 1 |
| *Gna11* | 1 |
| *Kcnk6* | 2 |
| *Fkbp1b* | 3 |
| *Pla2g5* | 3 |
| *F2rl1* | 2 |
| *Ptger1* | 3 |
| *Adora3* | 2 |
| *Gnaq* | 2 |
| *Wnk3* | 2 |
| *Lyz2* | 4 |
| *Agxt2* | 2 |
| *Fga* | 4 |
| *Mybph* | 1 |
| *Nox1* | 5 |
| *Ptger2* | 3 |
| *Asl* | 2 |

### C. Gene-Gene and Gene-Phenotype Embedding Spaces Achieved Higher Gene Classification Performance Than Phenotype-Gene Embedding Spaces

Since we utilized multiple gene and phenotype datasets, we analyzed the contribution of single embedding spaces to the classification results across different network combinations (Fig. 2). We observed that the Gene-Gene embeddings performed better than the Gene-Phenotype embeddings when the gene network was multiplex. When the gene network was aggregated, the Gene-Phenotype embeddings performed significantly better than the Gene-Gene embeddings, whereas when the phenotype network was aggregated, Gene-Gene embeddings performed significantly better than the Gene-Phenotype embeddings. On the other hand, there was no significant difference between their performances when both gene and phenotype networks were aggregated. Moreover, the results clearly showed that the Gene-Gene and Gene-Phenotype embedding spaces performed significantly higher than the Phenotype-Gene embeddings for all network combinations.

**Fig. 2.**
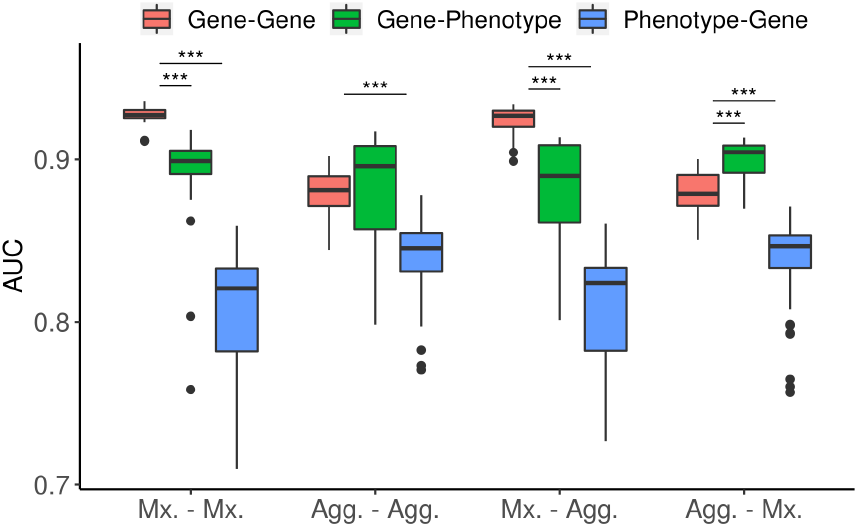
Contribution of gene-gene, gene-phenotype and phenotype-gene embedding spaces across different network combinations which utilize all three multiplex (Mx.) or aggregated (Agg.) gene and phenotype networks. Groups are shown in Gene-Phenotype network format. ***: *p* ≤ 0.001

### D. Multiplex Gene Networks Achieved Higher Gene Classification Performance Than Aggregated Gene Networks

We investigated the effect of multiplex networks of genes and phenotypes on the classification performance by using all available layers and omitting single gene/phenotype layers (Fig. 3 and Fig. 4). Here, we did not include gene embeddings from Phenotype-Gene neighborhood space, as their contribution to the disease-gene classification performance was very little and not consistent. First, we compared the classification performance when the phenotype network was multiplex and gene network was multiplex or aggregated. We observed that the classification performance with multiplex gene networks was significantly higher than the performance with aggregated gene networks when the Gene-Gene embeddings were utilized (Fig. 3. A, B, C, D). Except for a significant but modest difference in Fig. 3.A, for Gene-Phenotype embeddings, we did not observe a difference when the multiplex gene network was used. Overall, when we concatenated Gene-Gene and Gene-Phenotype embeddings, we observed higher AUC scores for multiplex gene networks than aggregated gene networks (Supplementary Table 4). The range of mean AUC for the four multiplex gene networks in Fig. 3 was [92.69% - 94.85%] when phenotype network was multiplex of three layers, while the mean AUC range for aggregated gene networks was [90.71% - 93.56%] when gene network was multiplex of three layers.

**Fig. 3.**
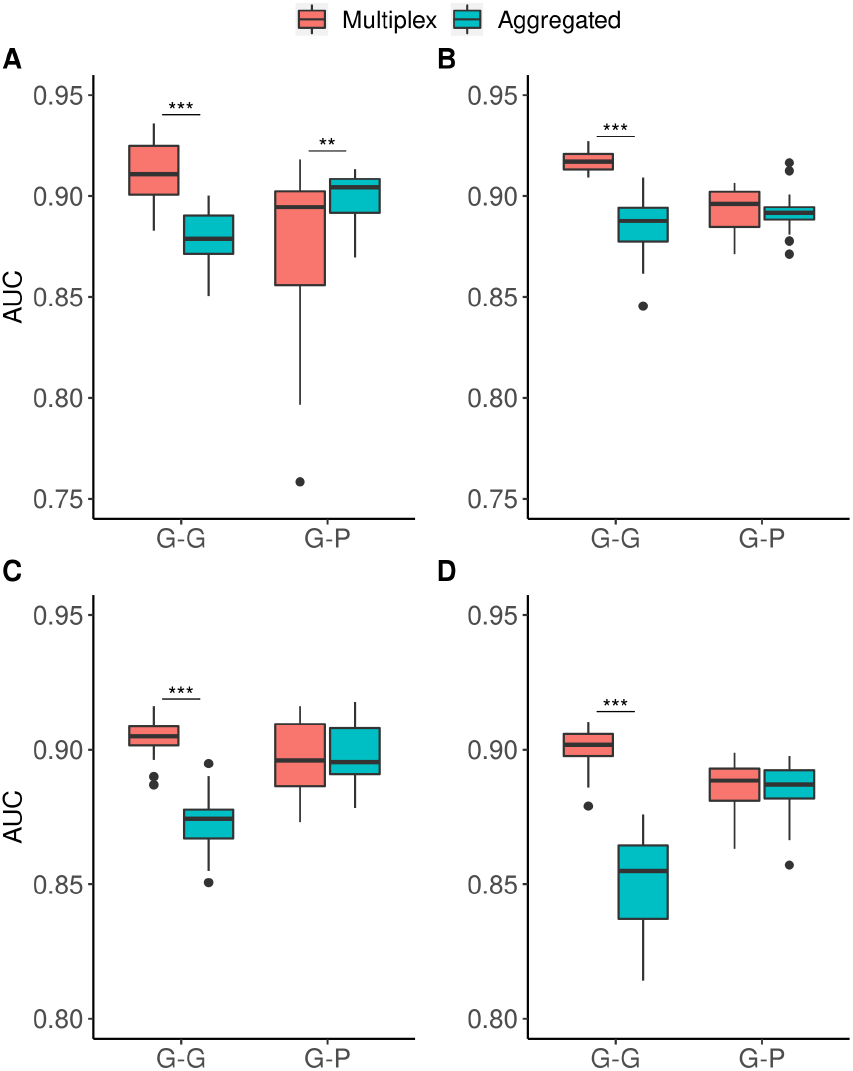
Effect of multiplex vs aggregated gene networks on the gene classification performance. All eight networks have multiplex networks of three phenotype layers and multiplex vs aggregated layers of A) (PPI, PWY, CO-EXPR), B) (PPI, PWY), C) (PPI, CO-EXPR), and D) (PWY, CO-EXPR). PWY: Pathway, CO-EXPR: Co-expression. **: *p* ≤ 0.01, ***: *p* ≤ 0.001

**Fig. 4.**
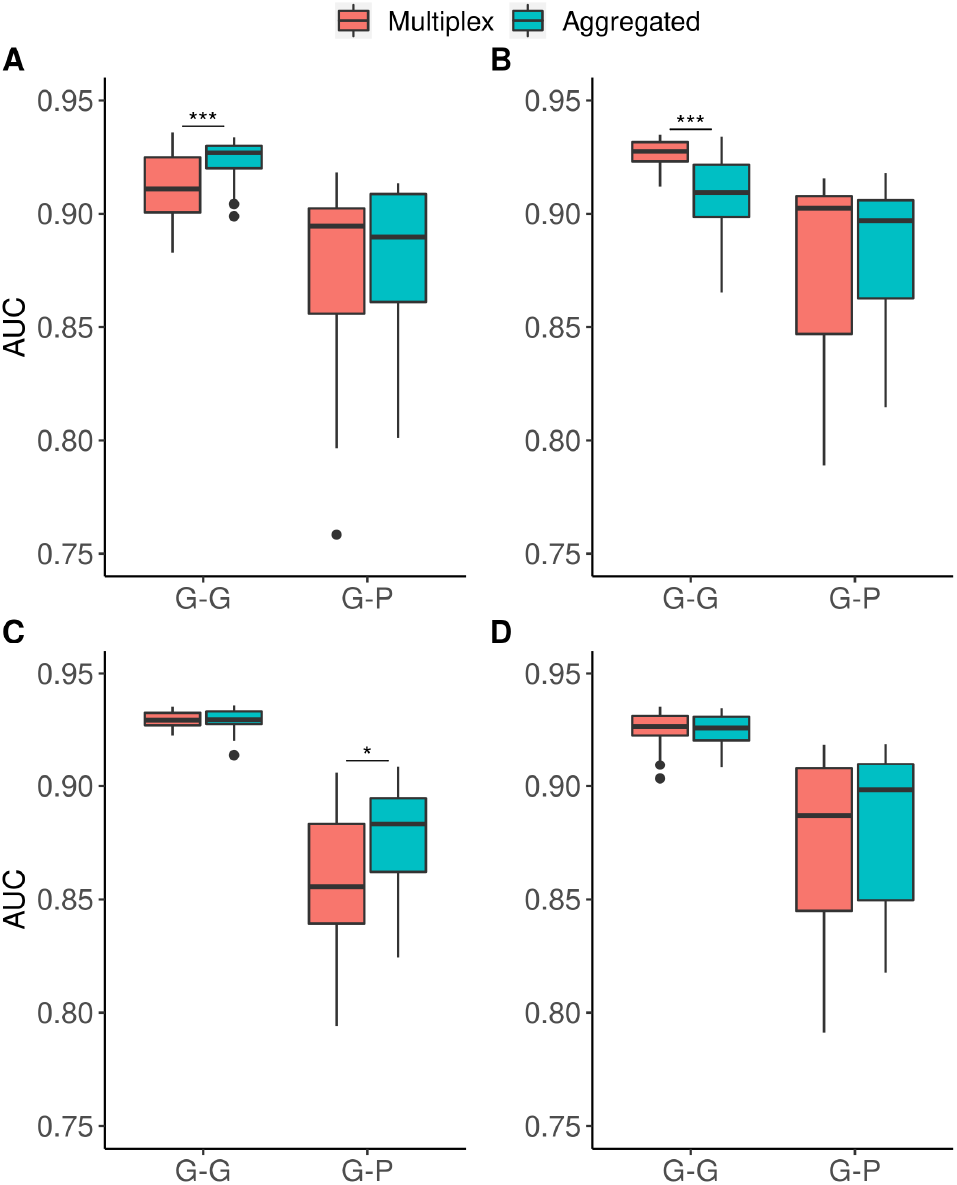
Effect of multiplex vs aggregated phenotype networks on the gene classification performance. All eight networks have multiplex network of three gene layers and multiplex vs. aggregated layers of A) (MPO, DO, QP), B) (MPO, QP), C) (DO, QP), D) (MPO, DO). *: *p* ≤ 0.05, ***: *p* ≤ 0.001

Next, we compared the classification performance when the gene network was multiplex, and the phenotype network was multiplex or aggregated. We did not observe a similar trend for multiplex phenotype networks compared to aggregated phenotype networks. There was no statistically significant difference between Gene-Phenotype embeddings’ performances on three network configurations (Fig. 4. A, B, D). For one configuration, Gene-Phenotype embeddings of aggregated network achieved higher score than the multiplex network (Fig. 4. C). Gene-Gene embeddings showed mixed results, for one case Gene-Gene embedding for aggregated phenotype network performed better (Fig. 4. A), for another case Gene-Gene embedding for multiplex phenotype network performed better (Fig. 4. B), and for others there was not statistically significant difference (Fig. 4. C-D).

### E. PPI Layer has the Largest Contribution to the Classification Performance

We analyzed the effects of the gene and phenotype layers to evaluate their individual contributions to the gene classifications. When we omitted the PPI layer from multiplex gene-phenotype networks, the classification AUC for the concatenated Gene-Gene and Gene-Phenotype embeddings dropped to 92.69% (±0.38%) from 94.85% (±0.29%). Omitting the co-expression layer dropped the AUC to 94.42% (±0.30%) and omitting the pathway layer dropped the AUC to 93.99% (±0.44%), suggesting the contribution from the co-expression layer was the smallest. We observed a similar trend for the single embedding space classification performances (Fig. 3). We observed a drop in AUC for Gene-Phenotype embeddings when we omitted one phenotype layer from the reference multiplex network, which had three phenotype layers (Fig. 4). But surprisingly we observed an increase in AUC for single Gene-Gene embeddings.

### F. Hyperparameter Analysis

We investigated the effects of *top N*, *embedding size* and *context size* parameters of NECo on the gene disease classification. NECo generates the embeddings separately with different hyperparameters, and then those embeddings are concatenated and used in a supervised/unsupervised algorithm. Therefore, we performed hyperparameter analysis for each neighborhood embeddings separately. We studied the parameter sensitivity of NECo as measured by the classification performance determined by AUC. We utilized a multiplex three-layer gene and three-layer phenotype network structure. Supplementary Fig. 1 shows the classification performance as a function of each of the three parameters when fixing the other two for three different neighborhoods, namely Gene-Gene (Supplementary Fig. 1. A, D, G), Gene-Phenotype (Supplementary Fig. 1. B, E, H) and Phenotype-Gene (Supplementary Fig. 1. C, F, I). We fixed the *embedding size* at 150 for all neighborhoods and set the default value of *context size* to 5, 50 and *top N* for Gene-Gene, Phenotype-Gene, and Gene-Phenotype neighborhoods, respectively as they had better performance around these *context size* values.

We observed that gene classification performance was consistently high for *top* N ≤ 350 when Gene-Gene and Gene-Phenotype embedding spaces were utilized (Supplementary Fig. 1. A-B). On the other hand, for the Phenotype-Genotype embedding space there seemed to be a variation in classification performance and classification performance peaked for *top N* = 500 (Supplementary Fig. 1.C).

We observed that the classification performance had a peak value for the Gene-Gene embedding space when the *context size* = 5 (Supplementary Fig. 1.D) and dropped for larger values of *context size*. For the Gene-Phenotype neighborhood, as we had one gene in the neighborhood, the performance peaked when the *context size* converged to *top N* (Supplementary Fig. 1.E). As Gene-Phenotype neighborhood consists of one gene at the beginning of neighborhood and the rest consists of phenotypes, to learn gene embeddings using the Gene-Phenotype neighborhood requires the use of larger *context sizes* for the Skip-gram algorithm. The classification performance starts to converge when *context size* = 75 for the Phenotype-Gene neighborhood (Supplementary Fig. 1.F).

We observed that different Gene-Gene *embedding sizes* did not have much impact on the classification performance (Supplementary Fig. 1.G). For the Gene-Phenotype neighborhood space, embedding size of 200 and larger had high classification performance consistently (Supplementary Fig. 1.H). For the Phenotype-Gene embedding space, *embedding size* = 150 had the top performance, but there were fluctuations in the performance scores (Supplementary Fig. 1.I). These results suggest that it is easy to find better performing hyperparameter values for NECo especially for Gene-Gene and Gene-Phenotype embedding spaces, but for the Phenotype-Genotype embedding space there is a variation in classification performance with respect to *embedding size*.

## IV. Discussion

In this study, we developed a node embedding tool called NECo that can utilize multiplex heterogeneous networks of genes and phenotypes. NECo uses stationary node ranks of RWR as a proximity measure of nodes, divides the node ranks into different neighborhood spaces, and then applies the Skipgram algorithm to generate the node embeddings.

To evaluate the performance of NECo, we applied it to multiple gene- and phenotype-related rat datasets to classify hypertension disease-related genes. We compared NECo to the state-of-the-art node embedding methods DeepWalk, Node2vec and Metapath2vec by determining how accurately they identified a known set of 167 hypertension-related genes. NECo achieved a 94.97% AUC and outperformed the closest performer by about 9% margin. In addition, NECo was able to identify hypertension-related genes not previously annotated in RGD. As we showed, 18 of the 20 top ranked novel predicted genes, based on NECo embeddings, were found to have literature that supported their association with hypertension. The ranked list of novel predictions can be further analyzed to identify possible pathways and mechanisms of hypertension. For example, we performed GO enrichment analysis of the 100 top ranked genes using the MOET tool at RGD. Interestingly, a cluster of over 21 of the 100 genes were enriched for the GO:Biological Process Ontology term ‘blood circulation’ (GO:0008015; Bonferroni corrected p 1.29E-18). Furthermore, a cluster of 10 of those 21 genes showed enrichment in GO:Molecular Function Ontology term ‘G protein-coupled receptor activity’ (GO: 0004930; Bonferroni corrected p 1.70E-7). These data identify putative novel candidate hypertension genes and implicate dysregulation in G protein-coupled receptor signaling as a candidate mechanism. Similarly, NECo could be applied to other complex diseases by integrating multidimensional molecular and physiological data to identify novel genes and pathways.

We observed that, use of gene embeddings from Phenotype-Gene neighborhood space did not contribute to the classification performance of gene embeddings. This could be resulted because of the similarity of gene-phenotype and phenotype-gene neighborhoods as we expect RWR to generate symmetric rankings to some extent for these neigborhoods.

NECo’s power of using multiplex networks could be utilized more efficiently if the two types of node embeddings were used in the downstream analysis. We did not generate phenotype embeddings for the current experimental study, but it is straightforward to generate phenotype embeddings for a different problem setting. A higher contribution of gene multiplex networks to the gene embeddings in the gene classification task compared to the contribution of multiplex phenotype networks suggests that if the embeddings of both node types are used in the downstream analysis, the power of multiplex network usage for both node types would be efficiently utilized in the downstream analysis such as drug-gene interaction studies or gene/cell line classification. Furthermore, NECo can be compared to more recent multiplex heterogeneous node embedding algorithms for different type of downstream analyses.

## Acknowledgment

This manuscript is dedicated to the memory of Prof. M. Shimoyama who passed away on Feb. 19, 2020. The authors thanks to Dr. Arjun Krishnan for fruitful discussions.

## Notes

This study was funded by grant R01 HL064541 from the National Heart, Lung, and Blood Institute of the National Institutes of Health (RGD) and by grant R35GM133657 from National Institute of General Medical Sciences of the National Institutes of Health.

### Competing Interest Statement

The authors have declared no competing interest.

### Summary of Updates

Section III.C clarified, supplemental information updated.

