## Supplementary Information for "NECo: A node embedding algorithm for multiplex heterogeneous networks"

Jennifer R. Smith, G. Thomas Hayman, Anne E. Kwitek
*Rat Genome Database*
*Department of Biomedical Engineering*
*Department of Physiology*
*Medical College of Wisconsin*
Milwaukee WI USA

Serdar Bozdag
Department of Computer Science
Marquette University
Milwaukee WI USA


**Supplementary Table 1**. Hypertension Disease Related Mammalian Phenotype Ontology Terms

| TERM ID | TERM |
|---|---|
| MP:0000230 | abnormal systemic arterial blood pressure |
| MP:0000231 | hypertension |
| MP:0002842 | increased systemic arterial blood pressure |
| MP:0003034 | increased pulmonary vascular resistance |
| MP:0003328 | portal hypertension |
| MP:0003349 | abnormal circulating renin level |
| MP:0003352 | increased circulating renin level |
| MP:0003547 | abnormal pulmonary pressure |
| MP:0003548 | pulmonary hypertension |
| MP:0003819 | increased left ventricle diastolic pressure |
| MP:0003820 | increased left ventricle systolic pressure |
| MP:0003823 | increased left ventricle developed pressure |
| MP:0004012 | increased pulmonary artery pressure |
| MP:0004118 | abnormal baroreceptor morphology |
| MP:0004164 | abnormal neurohypophysis morphology |
| MP:0004167 | abnormal cingulate gyrus morphology |
| MP:0004184 | abnormal baroreceptor physiology |
| MP:0004216 | salt-resistant hypertension |
| MP:0004217 | salt-sensitive hypertension |
| MP:0004875 | increased mean systemic arterial blood pressure |
| MP:0004878 | increased systemic vascular resistance |
| MP:0004879 | decreased systemic vascular resistance |
| MP:0005522 | increased circulating atrial natriuretic factor |
| MP:0005529 | abnormal renal vascular resistance |
| MP:0005530 | decreased renal vascular resistance |
| MP:0005531 | increased renal vascular resistance |
| MP:0005532 | abnormal vascular resistance |
| MP:0005611 | decreased circulating antidiuretic hormone level |
| MP:0006143 | increased systemic arterial diastolic blood pressure |
| MP:0006144 | increased systemic arterial systolic blood pressure |
| MP:0006265 | increased pulse pressure |
| MP:0006373 | abnormal circulating angiotensinogen level |
| MP:0006375 | increased circulating angiotensinogen level |
| MP:0008776 | increased right ventricle peak pressure |
| MP:0008777 | increased right ventricle diastolic pressure |
| MP:0010695 | abnormal blood pressure regulation |
| MP:0010697 | abnormal systemic arterial blood pressure regulation |
| MP:0010758 | increased right ventricle systolic pressure |
| MP:0011022 | abnormal circadian regulation of systemic arterial blood pressure |
| MP:0011312 | abnormal kidney afferent arteriole morphology |

| | |
|---|---|
| MP:0012007 | abnormal chloride level |
| MP:0012045 | increased susceptibility to hypertension |
| MP:0012046 | decreased susceptibility to hypertension |

**Supplementary Table 2.** Hypertension Disease Related Disease Ontology Terms

| TERM ID | TERM |
|---|---|
| DOID:1073 | renal hypertension |
| DOID:10763 | hypertension |
| DOID:10824 | malignant hypertension |
| DOID:6432 | pulmonary hypertension |
| DOID:9003234 | Hypertensive Nephropathy |
| DOID:9006166 | Pulmonary Hypertension, Hypoxia-Induced |

**Supplementary Table 3.** Hypertension disease related genes having experimental annotations at Rat Genome Database. The genes having only expression based experimental evidence codes are excluded. IDA: Inferred from direct assay; IAGP: Inferred from association of genotype from phenotype; IMP: Inferred from mutant phenotype.

| RGD ID | Gene Symbol | Disease Ontology Term | Evidence |
|---|---|---|---|
| 2031 | Ada | hypertension | IDA |
| 2041 | Add1 | hypertension | IAGP |
| 2041 | Add1 | hypertension | IDA |
| 2042 | Add2 | hypertension | IDA |
| 2043 | Add3 | hypertension | IDA |
| 2048 | Adora1 | hypertension | IDA |
| 2048 | Adora1 | hypertension | IMP |
| 2057 | Adra2b | hypertension | IMP |
| 2059 | Adrb1 | renovascular hypertension | IDA |
| 2059 | Adrb1 | renovascular hypertension | IMP |
| 2069 | Agt | hypertension | IDA |
| 2069 | Agt | hypertension | IMP |
| 2069 | Agt | pulmonary hypertension | IDA |
| 2070 | Agtr1a | Hypertensive Nephropathy | IMP |
| 2070 | Agtr1a | hypertension | IMP |
| 2070 | Agtr1a | renovascular hypertension | IMP |
| 2072 | Agtr2 | hypertension | IMP |
| 2072 | Agtr2 | renovascular hypertension | IDA |
| 2072 | Agtr2 | renovascular hypertension | IMP |
| 2081 | Akt1 | hypertension | IDA |
| 2096 | Alox5 | hypertension | IDA |
| 2096 | Alox5 | pulmonary hypertension | IDA |
| 2096 | Alox5 | pulmonary hypertension | IMP |
| 2097 | Alox5ap | pulmonary hypertension | IMP |
| 2147 | Ar | hypertension | IDA |
| 2147 | Ar | hypertension | IMP |
| 2150 | Arg1 | hypertension | IMP |
| 2151 | Arg2 | hypertension | IMP |
| 2167 | Atp1a1 | hypertension | IAGP |
| 2186 | Avpr2 | hypertension | IDA |
| 2200 | Bcl2l1 | pulmonary hypertension | IDA |
| 2201 | Bdkrb2 | pulmonary hypertension | IDA |
| 2232 | C3 | pre-eclampsia | IDA |
| 2275 | Casp3 | hypertension | IDA |
| 2279 | Cat | Renoprival Hypertension | IMP |
| 2369 | Cnr1 | hypertension | IMP |
| 2379 | Comt | hypertension | IDA |
| 2453 | Cyp11b1 | hypertension | IAGP |
| 2454 | Cyp11b2 | hypertension | IMP |

| | | | |
|---|---|---|---|
| 2493 | Ace | Hypertensive Nephrosclerosis | IMP |
| 2493 | Ace | hypertension | IDA |
| 2493 | Ace | hypertension | IMP |
| 2493 | Ace | renovascular hypertension | IDA |
| 2503 | Nqo1 | hypertension | IDA |
| 2521 | Drd3 | hypertension | IDA |
| 2535 | Ednra | hypertension | IDA |
| 2535 | Ednra | pulmonary hypertension | IDA |
| 2535 | Ednra | pulmonary hypertension | IMP |
| 2536 | Ednrb | hypertension | IMP |
| 2536 | Ednrb | pulmonary hypertension | IDA |
| 2536 | Ednrb | pulmonary hypertension | IMP |
| 2582 | Esr2 | hypertension | IMP |
| 2582 | Esr2 | pulmonary hypertension | IDA |
| 2586 | F2r | hypertension | IDA |
| 2609 | Fgf2 | pulmonary hypertension | IMP |
| 2624 | Fn1 | renovascular hypertension | IDA |
| 2645 | G6pd | Pulmonary Hypertension, Hypoxia-Induced | IMP |
| 2692 | Gja5 | hypertension | IDA |
| 2703 | Glp1r | hypertension | IDA |
| 2729 | Gpx1 | hypertension | IDA |
| 2741 | Nr3c1 | hypertension | IMP |
| 2802 | Hmgb1 | pulmonary hypertension | IMP |
| 2803 | Hmgcr | pulmonary hypertension | IMP |
| 2806 | Hmox1 | hypertension | IDA |
| 2806 | Hmox1 | pulmonary hypertension | IDA |
| 2806 | Hmox1 | renal hypertension | IDA |
| 2857 | Icam1 | pre-eclampsia | IMP |
| 2858 | Id1 | pulmonary hypertension | IDA |
| 2868 | Igf1 | hypertension | IAGP |
| 2891 | Il1b | hypertension | IDA |
| 2901 | Il6 | eclampsia | IDA |
| 2901 | Il6 | pulmonary hypertension | IDA |
| 2916 | Ins2 | hypertension | IDA |
| 2943 | Jun | hypertension | IDA |
| 2953 | Kcna5 | pulmonary hypertension | IDA |
| 2960 | Kcnj8 | hypertension | IDA |
| 2961 | Kcnmb1 | hypertension | IDA |
| 3000 | Lep | hypertension | IDA |
| 3015 | Lox | hypertension | IDA |
| 3030 | Smad1 | pulmonary hypertension | IDA |
| 3049 | Mas1 | hypertension | IMP |
| 3098 | Mme | hypertension | IMP |

| | | | |
|---|---|---|---|
| 3100 | Mmp7 | hypertension | IMP |
| 3130 | Myc | pulmonary hypertension | IDA |
| 3175 | Klk1b3 | hypertension | IAGP |
| 3177 | Ngfr | hypertension | IAGP |
| 3184 | Nos1 | hypertension | IDA |
| 3184 | Nos1 | hypertension | IMP |
| 3185 | Nos2 | renal hypertension | IMP |
| 3186 | Nos3 | hypertension | IDA |
| 3186 | Nos3 | pulmonary hypertension | IDA |
| 3186 | Nos3 | pulmonary hypertension | IMP |
| 3186 | Nos3 | renal hypertension | IDA |
| 3193 | Nppa | pulmonary hypertension | IDA |
| 3194 | Nppb | hypertension | IDA |
| 3195 | Npr1 | hypertension | IAGP |
| 3197 | Npy | hypertension | IMP |
| 3198 | Npy1r | hypertension | IMP |
| 3238 | Oxt | hypertension | IMP |
| 3249 | Serpine1 | hypertension | IDA |
| 3249 | Serpine1 | pulmonary hypertension | IDA |
| 3326 | Serpina1 | pulmonary hypertension | IDA |
| 3329 | Pik3r1 | hypertension | IMP |
| 3346 | Plcd1 | hypertension | IDA |
| 3369 | Ppara | Hypertensive Nephropathy | IDA |
| 3369 | Ppara | hypertension | IMP |
| 3371 | Pparg | hypertension | IDA |
| 3371 | Pparg | hypertension | IMP |
| 3395 | Prkca | hypertension | IAGP |
| 3396 | Prkcb | hypertension | IDA |
| 3438 | Ptgis | pulmonary hypertension | IDA |
| 3438 | Ptgis | pulmonary hypertension | IMP |
| 3439 | Ptgs1 | renovascular hypertension | IMP |
| 3443 | Ptk2 | hypertension | IAGP |
| 3454 | Ptprj | hypertension | IAGP |
| 3554 | Ren | hypertension | IDA |
| 3555 | Resp18 | hypertension | IMP |
| 3645 | Ccl2 | hypertension | IDA |
| 3714 | Slc6a4 | pulmonary hypertension | IDA |
| 3714 | Slc6a4 | pulmonary hypertension | IMP |
| 3720 | Slc9a3 | hypertension | IDA |
| 3732 | Sod2 | hypertension | IDA |
| 3733 | Sod3 | hypertension | IDA |
| 3752 | Spp1 | renovascular hypertension | IDA |
| 3807 | Tac1 | pulmonary hypertension | IMP |

| | | | |
|---|---|---|---|
| 3810 | Tacr3 | hypertension | IDA |
| 3811 | Tacr1 | pulmonary hypertension | IMP |
| 3812 | Tacr2 | pulmonary hypertension | IMP |
| 3826 | Tbxas1 | hypertension | IDA |
| 3852 | Tgfbr1 | pulmonary hypertension | IMP |
| 3876 | Tnf | Hypertension, Pregnancy-Induced | IMP |
| 3876 | Tnf | hypertension | IAGP |
| 3876 | Tnf | hypertension | IMP |
| 3903 | Trh | hypertension | IDA |
| 3904 | Trhr | hypertension | IDA |
| 3930 | Uts2 | hypertension | IDA |
| 3932 | Ucp2 | hypertension | IDA |
| 61276 | Crhr1 | hypertension | IDA |
| 61925 | Prkce | hypertension | IAGP |
| 61927 | Pecam1 | pulmonary hypertension | IDA |
| 62043 | Xdh | hypertension | IDA |
| 62086 | Acsm3 | hypertension | IAGP |
| 67383 | Prkcd | hypertension | IAGP |
| 68371 | Mtor | hypertension | IDA |
| 68371 | Mtor | pulmonary hypertension | IMP |
| 68407 | Lrp2 | hypertension | IDA |
| 68411 | Sh2b3 | hypertension | IMP |
| 69305 | Nr2f2 | hypertension | IMP |
| 70499 | Birc5 | pulmonary hypertension | IMP |
| 71061 | Kynu | hypertension | IAGP |
| 619754 | Smpd3 | pulmonary hypertension | IMP |
| 619790 | Rag1 | Hypertensive Nephropathy | IMP |
| 619790 | Rag1 | hypertension | IMP |
| 619796 | Cysltr1 | pulmonary hypertension | IMP |
| 619806 | Mtpn | hypertension | IDA |
| 619830 | Cd40 | pulmonary hypertension | IMP |
| 619866 | Dynll1 | hypertension | IMP |
| 619993 | Atp5f1a | pulmonary hypertension | IDA |
| 620007 | Cyp2j4 | pre-eclampsia | IMP |
| 620209 | Cxcl10 | renovascular hypertension | IDA |
| 620217 | Slc5a2 | hypertension | IDA |
| 620243 | Gnai2 | hypertension | IMP |
| 620293 | Ece1 | hypertension | IDA |
| 620293 | Ece1 | pulmonary hypertension | IMP |
| 620322 | Cldn16 | hypertension | IDA |
| 620349 | Ptgs2 | hypertension | IAGP |
| 620349 | Ptgs2 | pulmonary hypertension | IMP |
| 620396 | Kl | hypertension | IDA |

| | | | |
|---|---|---|---|
| 620401 | Bdkrb1 | hypertension | IMP |
| 620446 | Shc1 | Hypertensive Nephropathy | IMP |
| 620573 | Cyba | hypertension | IDA |
| 620573 | Cyba | pulmonary hypertension | IDA |
| 620600 | Nox4 | hypertension | IMP |
| 620630 | Hrh3 | hypertension | IDA |
| 620713 | Fgfr1 | pulmonary hypertension | IMP |
| 620732 | Ephx2 | hypertension | IDA |
| 620732 | Ephx2 | hypertension | IMP |
| 620732 | Ephx2 | pulmonary hypertension | IMP |
| 620795 | Src | hypertension | IDA |
| 620809 | Slc12a2 | hypertension | IDA |
| 620948 | Abat | hypertension | IDA |
| 620995 | Pde5a | pre-eclampsia | IMP |
| 620995 | Pde5a | pulmonary hypertension | IMP |
| 621057 | Tnc | pulmonary hypertension | IDA |
| 621137 | Rnpep | hypertension | IMP |
| 621159 | Il1rn | hypertension | IDA |
| 621163 | Mif | hypertension | IDA |
| 621316 | Mmp2 | renal hypertension | IDA |
| 621320 | Mmp9 | pulmonary hypertension | IDA |
| 621320 | Mmp9 | renal hypertension | IDA |
| 621376 | Atp5pf | pulmonary hypertension | IMP |
| 621503 | Kcnq1 | hypertension | IAGP |
| 621506 | Mapk8 | hypertension | IDA |
| 621528 | Cxcr3 | renovascular hypertension | IDA |
| 621647 | Vip | pulmonary hypertension | IDA |
| 621862 | Atf2 | hypertension | IDA |
| 621884 | Uts2r | hypertension | IDA |
| 631375 | Egln1 | hypertension | IMP |
| 631401 | Wnk4 | hypertension | IAGP |
| 708418 | Cd40lg | pre-eclampsia | IMP |
| 1306950 | Nisch | hypertension | IMP |
| 1307690 | Nfatc2 | pulmonary hypertension | IMP |
| 1307917 | Mmp1 | pulmonary hypertension | IDA |
| 1310046 | Adamts16 | hypertension | IMP |
| 1310740 | Lipg | hypertension | IDA |
| 1559787 | Srf | hypertension | IDA |
| 1560646 | Myo6 | hypertension | IDA |
| 1563603 | Rad51 | hypertension | IDA |
| 1566119 | H2ax | hypertension | IDA |
| 1593188 | Eng | pre-eclampsia | IDA |

**Supplementary Table 4.** Mean AUC values with standard deviations are given for different combinations of multiplex and aggregated networks. The results are for 10 runs for each combination. PPI: protein-protein interaction, PWY: pathway, EXPR: co-expression, DO: Disease Ontology, MPO: Mammalian Phenotype Ontology, QP: Quantitative Phenotype Measurements

| | **Networks Used** | | **Mean AUC** | **Std. Dev.** |
|---|---|---|---|---|
| **Rank** | **Gene** | **Phenotype** | | |
| 1 | Multiplex (PPI, PWY, EXPR) | Aggregated (MPO, QP) | 94.97 | 0.24 |
| 2 | Multiplex (PPI, PWY, EXPR) | Aggregated (MPO, DO) | 94.92 | 0.25 |
| 3 | Multiplex (PPI, PWY, EXPR) | Aggregated (MPO, DO, QP) | 94.89 | 0.27 |
| 4 | Multiplex (PPI, PWY, EXPR) | Multiplex (MPO, DO) | 94.87 | 0.30 |
| 5 | Multiplex (PPI, PWY, EXPR) | Multiplex (MPO, QP) | 94.87 | 0.35 |
| 6 | Multiplex (PPI, PWY, EXPR) | Multiplex (MPO, DO, QP) | 94.85 | 0.29 |
| 7 | Multiplex (PPI, PWY, EXPR) | Multiplex (DO, QP) | 94.81 | 0.29 |
| 8 | Multiplex (PPI, PWY, EXPR) | Aggregated (DO, QP) | 94.52 | 0.12 |
| 9 | Multiplex (PPI, EXPR) | Multiplex (MPO, DO, QP) | 94.42 | 0.30 |
| 10 | Multiplex (PPI, PWY | Multiplex (MPO, DO, QP) | 93.99 | 0.44 |
| 11 | Aggregated (PPI, PWY, EXPR) | Multiplex (MPO, DO, QP) | 93.51 | 0.39 |
| 12 | Aggregated (PPI, PWY, EXPR) | Aggregated (MPO, DO, QP) | 93.29 | 0.34 |
| 13 | Multiplex (PWY, EXPR) | Multiplex (MPO, DO, QP) | 92.69 | 0.38 |

**Supplementary Table 5.** Novel gene predictions of NECo sorted by prediction probability.

| RGD ID | Symbol | Evidence |
|---|---|---|
| **3277** | Pdc | [2, 10, 26] |
| **3548** | Rcn2 | [19] |
| **619749** | Gna11 | [27] |
| **621450** | Kcnk6 | [21, 28] |
| **61835** | Fkbp1b | [6, 22, 40] |
| **62051** | Pla2g5 | [4, 23, 38] |
| **61977** | Ckmt2 | - |
| **620866** | F2rl1 | [20, 43] |
| **3434** | Ptger1 | [1, 24, 32] |
| **61935** | Pla2g10 | - |
| **2051** | Adora3 | [11, 44] |
| **620770** | Gnaq | [39, 43] |
| **1563131** | Wnk3 | [3, 12] |
| **3026** | Lyz2 | [17, 29, 30, 37] |
| **621767** | Agxt2 | [5, 13] |
| **2603** | Fga | [16, 31, 34, 35] |
| **620287** | Mybph | [8] |
| **620598** | Nox1 | [9, 18, 25, 33, 41] |
| **620020** | Ptger2 | [14, 36, 42] |
| **619974** | Asl | [7, 15] |

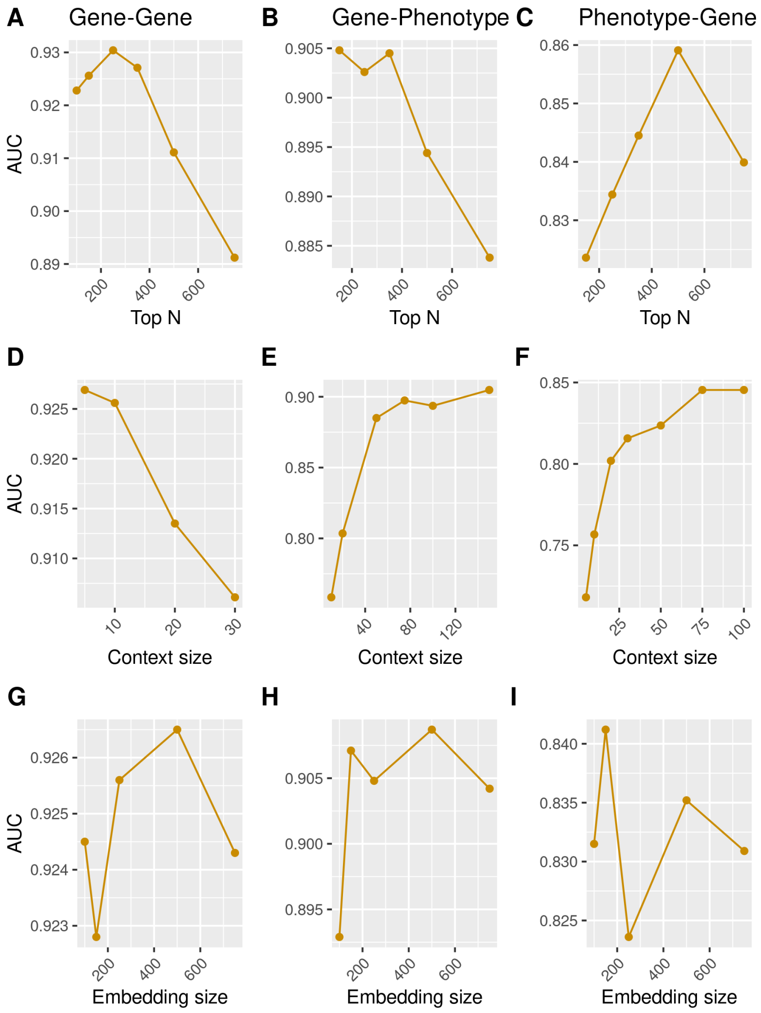


**Supplementary Fig. 1**. Parameter sensitivity of NECo. For each parameter evaluation the other two parameters are fixed. Panels by columns show the parameter plots by different embedding spaces.
